# Microbe-dependent heterosis in maize

**DOI:** 10.1101/2020.05.05.078766

**Authors:** Maggie R. Wagner, Clara Tang, Fernanda Salvato, Kayla M. Clouse, Alexandria Bartlett, Shannon Sermons, Mark Hoffmann, Peter J. Balint-Kurti, Manuel Kleiner

## Abstract

Hybrids account for nearly all commercially planted varieties of maize and many other crop plants, because crosses between inbred lines of these species produce F1 offspring that greatly outperform their parents. The mechanisms underlying this phenomenon, called *heterosis* or hybrid vigor, are not well understood despite over a century of intensive research (1). The leading hypotheses—which focus on quantitative genetic mechanisms (dominance, overdominance, and epistasis) and molecular mechanisms (gene dosage and transcriptional regulation)—have been able to explain some but not all of the observed patterns of heterosis (2, 3). However, possible ecological drivers of heterosis have largely been ignored. Here we show that heterosis of root biomass and germination in maize is strongly dependent on the belowground microbial environment. We found that, in some cases, inbred lines perform as well by these criteria as their F1 offspring under sterile conditions, but that heterosis can be restored by inoculation with a simple community of seven bacterial strains. We observed the same pattern for seedlings inoculated with autoclaved *vs.* live soil slurries in a growth chamber, and for plants grown in fumigated *vs.* untreated soil in the field. Together, our results demonstrate a novel, ecological mechanism for heterosis whereby soil microbes generally impair the germination and early growth of inbred but not hybrid maize.

## MAIN

In nature, all plants form close associations with diverse microbial symbionts that comprise a subset of the microbial species with which they share a habitat (4, 5). As part of their host’s environment, the host-associated microbial community (microbiome) can cause plasticity of important plant traits such as reproductive phenology, disease resistance, and general vigor (6–9). However, genetic variation within plant species affects not only microbiome assembly, but also the phenotypic response to microbes. Disentangling these relationships is a critical step toward understanding how plant-microbiome interactions evolved and how they can be harnessed for use in sustainable agriculture (10). Here, we describe our observation that perturbation of the soil microbial community disrupts *heterosis*, the strong and pervasive phenotypic superiority of hybrid maize genotypes relative to their inbred parent lines. To our knowledge this is the first report of microbial involvement in plant heterosis, a phenomenon of immense economic value and research interest.

In a previous field experiment, we observed that maize hybrids generally assemble rhizosphere microbiomes that are distinct from those of inbred lines. In addition, many microbiome features in F_1_ hybrids are not intermediate to those of their parent lines, suggesting that heterosis of plant traits is associated with heterosis of microbiome composition itself (11). To determine whether the same patterns manifest in a highly controlled environment where all microbial members are known, we developed gnotobiotic growth bags for growing individual maize plants in sterile conditions (see Methods). We planted surface-sterilized kernels of two inbred lines (B73 and Mo17) and their F_1_ hybrid (B73xMo17) in individual gnotobiotic growth bags containing autoclaved calcined clay hydrated with sterile 0.5x MS salt solution. The clay in each growth bag was inoculated with either a highly simplified synthetic community of seven bacterial strains known to colonize maize roots (12, 13) (~10^7^ CFU/mL) or a sterile buffer control. This system effectively eliminated contact between plants and external microbes (Supplementary Fig. 1b); however, it is possible that some kernels may have contained viable endophytes that could not be removed by surface-sterilization (Supplementary Fig. 1c). Genotypes and treatments were placed in randomized locations in a growth chamber under standard conditions (12-hr days, 27^℃^C/23^℃^C, ambient humidity). After four weeks, we harvested plants to investigate root colonization by these seven strains.

Unexpectedly, we observed that the inbred and hybrid plants were indistinguishable with respect to root and shoot fresh weight when grown in uninoculated growth bags, yet showed the expected heterotic pattern when grown with the synthetic bacterial community (Fig. 1a; Supplementary Table 1). This was due to a negative effect of the bacteria on both inbred genotypes rather than a positive effect on the hybrid. Although the synthetic community contained no known pathogens (13), it decreased the root weight of B73 and Mo17 seedlings by 48.4% [s.e.m. = 13.6%] and 60.8% [s.e.m. = 21.5%], respectively (Fig. 1). In contrast, the synthetic community reduced root weight of hybrids by only 19.2% [s.e.m. = 13.6%]. As a result, the strength of midparent heterosis was reduced from 100% in nonsterile conditions to 14% in sterile conditions (permutation test *P* = 0.002); a similar pattern was observed for shoot weight (*P* = 0.056; Fig. 1c-d). A separate experiment revealed that the synthetic community also lowered the germination rates for both inbred lines but not the hybrid (Supplementary Fig. 2). Germination of B73 after 4 days was 10.7% lower [s.e.m. = 4.3%] in the presence of the synthetic community relative to the sterile control; for Mo17, the synthetic community decreased germination rates by 32% [s.e.m. = 5.8%] (Supplementary Fig. 2).

**Figure 1.**
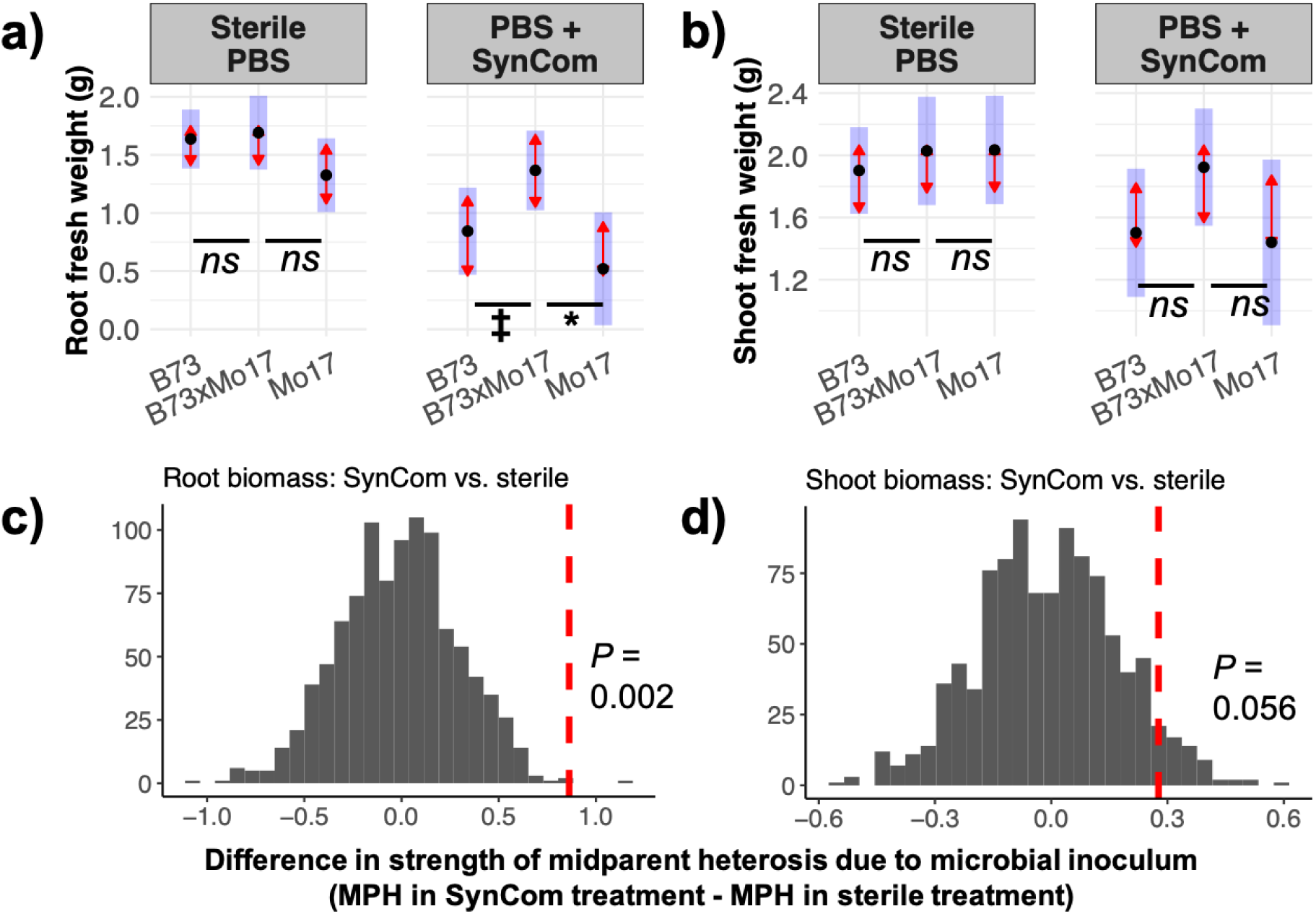
In Experiment 1, maize kernels were grown in calcined clay inoculated with phosphate-buffered saline (PBS) with or without 10^7^ CFU/mL of a synthetic community ("SynCom”) of seven bacterial strains. Sterile conditions reduced the strength of heterosis for **(a,c)** root biomass but not **(b,d)** shoot biomass. **(a-b)** Black points show the estimated marginal mean (EMM) trait values for each genotype in each treatment; blue rectangles show the 95% CIs for the EMMs. The red arrows show the 95% CIs for pairwise tests between genotypes in each treatment after correction for the family-wise error rate using Tukey’s procedure; non-overlapping arrows indicate statistically significant differences (alpha = 0.05). Detailed statistical results are provided in Supplementary Table 1. *N* = 14 per inbred genotype per treatment, *N =* 7 per hybrid genotype per treatment. ^***^ P<0.001; ^**^ P<0.01; ^*^ P<0.05; $ P<0.1; *ns* P>0.1 (Dunnett’s test of contrasts between each inbred line and the hybrid), **(c-d)** The strength of midparent heterosis (MPH) was calculated for each trait in each treatment using the EMM trait values. The observed difference in MPH between nonsterile and sterile treatments (Δ_MPH_) is shown as a vertical dashed line. The histogram shows the distribution of Δ_MPH_ for 999 permutations of the data with respect to treatment, *i.e.*, the distribution of Δ_MPH_ if there were no effect of treatment.

To determine whether natural, complex soil microbial communities also induce heterosis, we conducted a second growth chamber experiment with surface-sterilized kernels of the same three genotypes and a slightly modified protocol for gnotobiotic growth. We saturated the calcined clay medium in each growth bag with one of three treatments: a slurry derived from filtered farm soil, an autoclaved aliquot of the same slurry, or a sterile buffer control. Genotypes and treatments were arranged into randomized, replicated blocks in a growth chamber. We recorded the germination success or failure of each kernel and observed that the live soil slurry had a strong negative effect on germination of both inbred lines but not the hybrid (Fig. 2a). In the two sterile treatments, B73 and B73xMo17 germinated equally well. Mo17 still performed worse than B73xMo17, but the hybrid advantage was much less pronounced than it was in the live treatment. After one month we harvested all plants and measured fresh weights of roots and shoots. In growth bags that received the autoclaved slurry or sterile buffer treatments, all three genotypes produced root systems of equal biomass; in contrast, the hybrid’s root biomass was 18.3% higher than the midparent average when grown with the live soil slurry, consistent with the expected pattern of heterosis (Fig. 2b-c; Supplementary Table 2). Very poor germination of Mo17 prevented statistical comparison of its biomass to the hybrid in the live slurry treatment. Shoot biomass displayed the expected heterotic patterns with the hybrid out-performing the parental inbred lines under all conditions (Supplementary Fig. 3).

**Figure 2.**
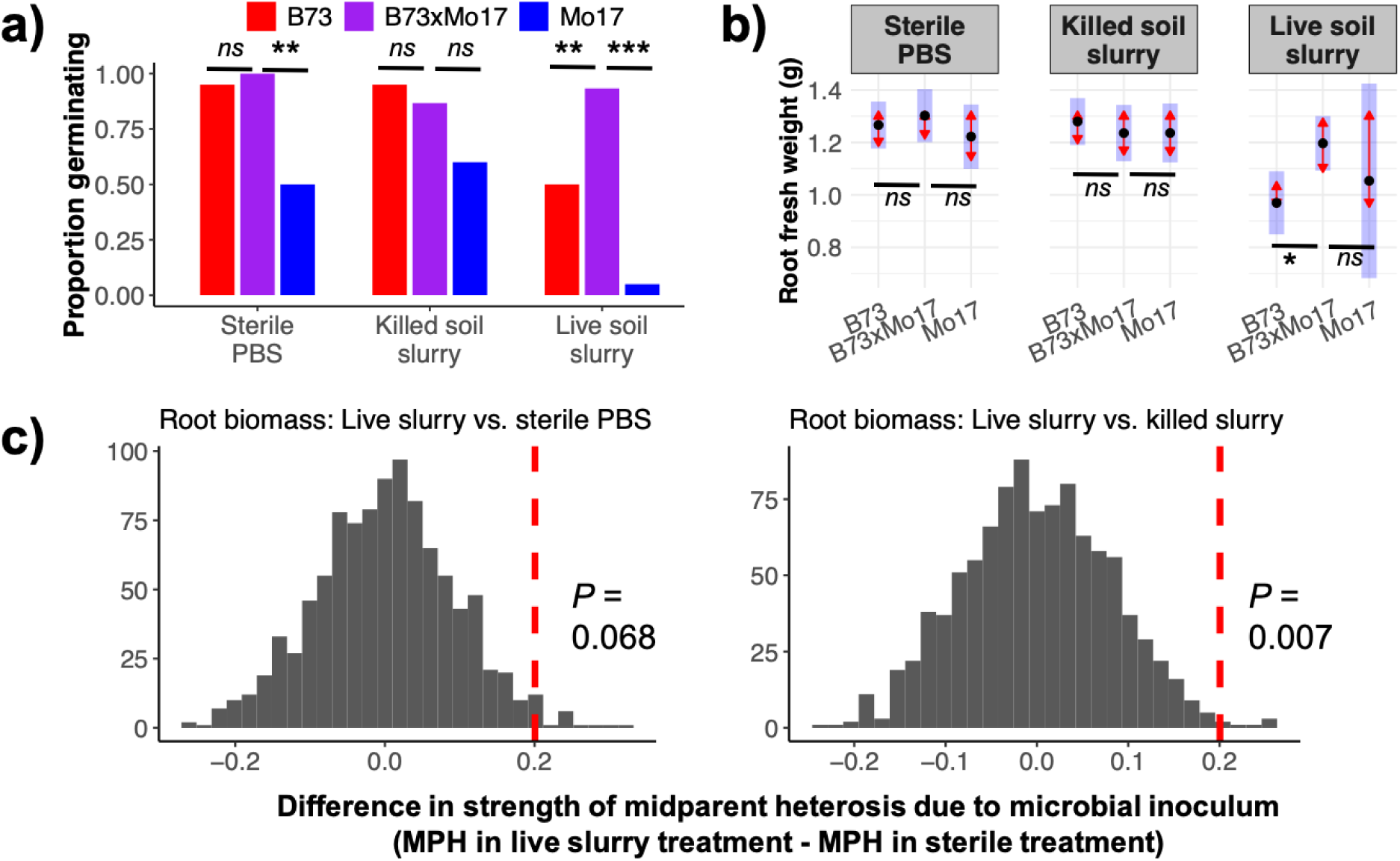
In Experiment 2, maize kernels were grown in calcined clay inoculated with sterile PBS, a live soil slurry in PBS, or an autoclaved (killed) aliquot of the same soil slurry. Sterile conditions reduced the strength of heterosis for germination success and root biomass. They did not affect heterosis of shoot biomass (Supplementary Figure 3). (a) Fisher’s Exact Test was used for statistical inference of germination proportions, (b) A linear mixed-effects model was used for statistical inference of root biomass. Black points show the estimated marginal mean (EMM) trait values for each genotype in each treatment; blue rectangles show the 95% CIs for the EMMs. The red arrows show the 95% CIs for pairwise tests between genotypes in each treatment after correction for the family-wise error rate using Tukey’s procedure; non-overlapping arrows indicate statistically significant differences (alpha = 0.05). Detailed statistical results are provided in Supplementary Table 2. ***N =*** 20 per inbred genotype per treatment, ***N =*** 15 per hybrid per treatment. ^***^ P<0.001; ^**^ P<0.01; ^*^ P<0.05; t F<0.1; ***ns*** P>0.1 (Dunnett’s test of contrasts between each inbred line and the hybrid), (c) The strength of midparent heterosis (MPH) was calculated for each treatment using the EMM trait values. The observed difference in MPH between nonsterile and sterile treatments (ΔMPH) is shown as a vertical dashed line. The histograms show the distributions of ΔMPH for 999 permutations of the data with respect to treatment, *i.e.*, the distribution of ΔMPH if there were no effect of treatment.

### Microbe-dependent heterosis in the field

Next, we conducted a field experiment to assess whether this phenomenon, which we termed “microbiota-dependent heterosis” or MDH, occurs in real soil under farm conditions. We planted surface-sterilized kernels of the same three genotypes into adjacent rows with four soil pre-treatments to perturb soil microbial community composition: (1) steamed, (2) fumigated with the mustard oil allyl isothiocyanate (AITC), (3) steamed and fumigated with AITC, (4) fumigated with chloropicrin, and (5) untreated control (Supplementary Fig. 4). All four treatments reduced the density of *Pythium* spp., a common phytopathogenic oomycete, relative to the untreated control (Supplementary Table 3); however, 2 weeks after treatment, counts of viable culturable bacteria were temporarily reduced only in the AITC + steam treatment, and only in shallow soil (Supplementary Fig. 5). Additionally, amplicon sequencing of the V4 region of the 16S rRNA gene and the fungal ITS1 confirmed that these treatments shifted the composition of the bacterial and fungal soil microbiomes relative to the control (Supplementary Fig. 6). The effects of the fumigation treatments persisted in the bulk soil for at least six weeks, and were also detected in the root microbiomes of juvenile plants at the end of the experiment (Supplementary Fig. 7). We monitored seedling emergence and measured leaf number and plant height at 15 days after planting (d.a.p.) and again at 27 d.a.p. After this final in-field measurement, we uprooted all plants in the control, chloropicrin, and AITC + steam treatments and measured their root and shoot biomass. Perturbation of the soil microbial community using chloropicrin or AITC + steam weakened heterosis of root biomass (Fig. 3; Supplementary Table 4). Additionally, all fumigation and steaming treatments decreased the strength of midparent heterosis for both height and leaf number (Supplementary Fig. 8). In contrast, heterosis of shoot dry weight was not affected. Rates of germination success did not differ consistently among treatments; although chloropicrin accelerated the germination of Mo17, the final germination proportions were similar among treatments (Supplementary Fig. 9). We note that AITC may influence plant development directly(14); however, the responses of each genotype to treatments involving AITC were generally congruent with responses to the non-AITC treatments.

**Figure 3.**
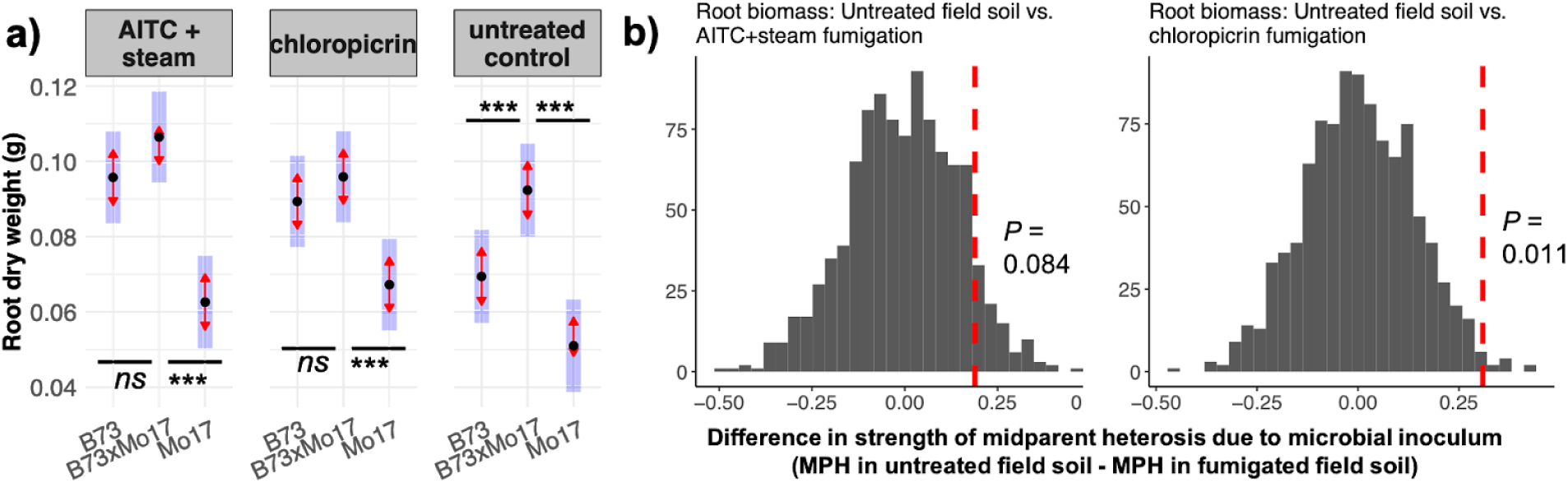
In Experiment 3, we grew maize in the field from seeds planted into untreated soil, soil fumigated with chloropicrin, or soil fumigated with AITC and steamed. Perturbation of soil microbiomes reduced heterosis for root biomass after 4 weeks of growth. Black points show the estimated marginal mean (EMM) trait values for each genotype in each treatment; blue rectangles show the 95% CIs for the EMMs. The red arrows show the 95% CIs for pairwise tests between genotypes in each treatment after correction for the family-wise error rate using Tukey’s procedure; non-overlapping arrows indicate statistically significant differences (alpha = 0.05). Detailed statistical results are provided in Supplementary Table 4. Effects on shoot biomass, plant height, and number of leaves are presented in Supplementary Figure 6. ***N =*** 56 per genotype per treatment. ^***^ **P<0.001; ^**^ P<0.01; ^*^ P<0.05; $ P<0.1; *ns* P>0.1** (Dunnett’s test of contrasts between each inbred line and the hybrid), **(b)** The strength of midparent heterosis (MPH) was calculated for each treatment using the EMM trait values. The observed difference in MPH between nonsterile and sterile treatments (ΔMPH) is shown as a vertical dashed line. The histograms show the distributions of ΔMPH for 999 permutations of the data with respect to treatment, i.e., the distribution of ΔMPH if there were no effect of treatment.

## Discussion

Our results suggest that interactions with soil-borne microbes are important for the expression of heterosis in maize. We observed microbe-dependent heterosis (MDH) in three independent experiments representing very different environmental contexts: in tightly controlled lab conditions with an inoculum of only seven bacterial strains (Fig. 1); in a growth chamber with a more complex microbial slurry derived from farm soil (Fig. 2); and in the field with or without soil fumigation (Fig. 3; Supplementary Fig. 8). This repeatability suggests that the mechanism could be quite general with respect to the causal microbes, although much more work would be needed to test the full range of natural soil microbiome diversity and the full range of plant genotypes.

In all of the cases presented above, MDH was driven not by beneficial microbes selectively boosting the performance of hybrids, but by soil-borne microbes selectively reducing the performance of inbred lines. This pattern is consistent with two possible, non-mutually-exclusive explanations. First, it may indicate that many or most soil microbes are weakly pathogenic to maize, and that hybrids are more resistant to them than are inbreds (the “Inbred Immunodeficiency hypothesis”). Second, it may reflect a costly defensive overreaction by inbreds, but not hybrids, to innocuous soil microbes (the “Inbred Immune Overreaction hypothesis”).

Multiple previous studies have described how plants that are immunocompromised through either genetic or chemical means can suffer infections that are not apparent in their immunocompetent neighbors. For example, maize mutants deficient in the defense hormone jasmonic acid were unable to grow to maturity in non-sterile soil in the field or greenhouse (15). Similarly, Arabidopsis mutant lines lacking three defense hormone signaling systems displayed reduced survival in wild soil (16). Application of glyphosate to bean plants temporarily arrested their growth in sterile soils; in non-sterile soils, however, the plants died quickly due to root infection by *Pythium* and *Fusarium* species (17). Because glyphosate inhibits the biosynthesis of phenylalanine and chorismite—which are precursors of several important components of the defense response including lignin, salicylic acid and phytoalexins—the study authors suggested that glyphosate predisposes the treated plants to infection by opportunistic pathogens to which they would otherwise be resistant (18). If weak pathogens drive MDH, then this implies that superior disease resistance in hybrids is a key mechanism of heterosis. Somewhat surprisingly, the effect of heterosis on plant disease resistance has not been well characterized. In maize, heterosis has been observed for resistance to anthracnose leaf blight and southern leaf blight but not to anthracnose stalk rot (19, 20). Heterosis for late blight resistance has also been noted in potato (21).

In contrast, the Inbred Immune Overreaction hypothesis does not require soil microbes to be pathogenic, but instead links MDH to the well-documented tradeoff between growth and genetic disease resistance (22). For instance, innocuous soil microbes could trigger a costly defensive response in inbreds but not in hybrid maize. The most detailed work on heterosis of disease resistance supports this hypothesis: in the model species Arabidopsis, hybrids displaying heterosis for growth and yield also displayed a *decreased* level of basal defense gene expression and *decreased* concentrations of the defense signaling hormone salicylic acid (23–26). However, despite their lower investment in constitutive defenses, the hybrids were not compromised in resistance to the biotrophic pathogen *Pseudomonas syringae*, nor in the inducible response to infection (26, 27).

All together, our results shed a new and unexpected light on the causes of heterosis, which have remained elusive despite over a century of investigation. They demonstrate the importance of ecological context for mapping genotype to phenotype, and generate new, testable hypotheses about the mechanisms of this widespread and critically important phenomenon. Many questions remain, and future work will require careful experimentation to delve into the molecular and physiological mechanisms of MDH and to assess the evidence for or against the Inbred Immunodeficiency hypothesis and the Inbred Immune Overreaction hypothesis. These new avenues of research have high potential to advance our understanding of heterosis in maize and many other crops, and to lead to new innovations for agricultural sustainability and productivity.

## METHODS

The genotypes used for experiments 1, 2, and 3 were B73, Mo17, and their F_1_ hybrid B73xMo17. All data analysis was performed using R version 3.5.3, particularly the packages lme4, tidyverse, lmerTest, car, and emmeans (28–32).

### Experiment 1

(December 2018). In a laminar flow hood, we placed kernels of each genotype into a sterile 7.5” x 15” Whirl-Pak self-standing bag (Nasco, Fort Atkinson, WI, USA) filled with 200 mL of autoclaved calcined clay (“Pro’s Choice Rapid Dry”; Oil-Dri Corporation, Chicago, IL). Immediately prior to planting, seeds were surface-sterilized using a 3-minute soak in 70% ethanol (v/v) followed by a 3-minute soak in 5% bleach (v/v) and three rinses with sterile deionized water; we plated extra seeds on malt extract agar (MEA) to confirm that this protocol was effective (Supplementary Fig. 1c). To each growth bag, we added 120 mL of either sterile 0.5x Murashige-Skoog basal salt solution (pH 6.0), or the same solution containing 10^7^ cells/mL of a synthetic community (SynCom) of seven bacterial strains known to colonize maize roots (12). We planted 28 kernels of each inbred line and 14 of the hybrid, divided evenly between the SynCom and control treatments, 4.5 cm deep using sterile forceps. The growth bags were sealed with sterile AeraSeal breathable film (Excel Scientific, Inc., Victorville, CA, USA) to allow gas exchange and then placed in randomized positions in a growth chamber (Percival Scientific Inc., Perry, IA). No additional liquid was added after the growth bags were sealed. After one month of growth (12-hr days, 27℃/23℃, ambient humidity), we opened the growth bags, uprooted the plants, rinsed off adhering clay, and patted them dry before measuring fresh weight of shoots and roots. We applied two-way ANOVA to linear models of biomass with Genotype, Treatment, and their interaction as predictor variables. *F*-tests with Type III sums of squares were used for significance testing, and pairwise contrasts were performed using Tukey’s *post-hoc* procedure.

### SynCom effects on germination

To test whether the SynCom affected germination, we conducted a 3×2×3 full factorial experiment manipulating plant genotype (B73, Mo17, and their F_1_ hybrid), microbial inoculant (SynCom *vs.* sterile control), and nutrient content (water, Hoagland’s solution, or MS). Five surface-sterilized kernels were placed onto filter paper in five petri dishes per genotype-inoculum-nutrient combination (*N* = 90 petri dishes) and inoculated with 2 mL of the SynCom (diluted to 10^6^ cells mL^−1^ in nutrient solution) or a sterile nutrient solution control. Petri dishes were incubated in the dark at 30℃ and germination rate was recorded for each dish after 4 days. We used the Kruskal-Wallis test for main effects of genotype, inoculum, and nutrient treatment and for an interaction between genotype and inoculum. Wilcoxon rank sum tests were used for pairwise contrasts; *P*-values were adjusted for multiple comparisons using the Benjamini-Hochberg false discovery rate (33).

### Experiment 2

(January 2019). To determine whether natural soil microbial communities produced the same effect as the SynCom, we used the same gnotobiotic growth bags as in Experiment 1 to compare plant growth in (1) a live soil slurry, (2) an autoclaved soil slurry, and (3) sterile buffer. We collected soil in November 2018 from field G4C at the Central Crops Research Station (Clayton, NC, USA) and stored it at 4^℃^C until use. We mixed 200 g of this soil into 1 L of phosphate-buffered saline (PBS) with 0.0001% Triton X-100 using a sterile spatula. The suspension was allowed to settle, filtered through Miracloth (22-25 µm pore size; Calbiochem, San Diego, CA, USA), and centrifuged for 30 minutes at 3,000 x *g*. The resulting pellet was resuspended in 200 mL of sterile PBS and immediately divided into two aliquots of 100 mL each. One aliquot was autoclaved for 30 minutes at 121^℃^C to produce a “killed” slurry concentrate. Live and killed soil slurry concentrates were diluted (10 mL slurry per L of 0.5x MS) to produce the final slurry treatments. An additional control consisted of diluted sterile PBS (10 mL PBS per L of 0.5x MS). Kernels were surface-sterilized as described above, planted in 150 mL sterile calcined clay, and hydrated with 90 mL of one of these three treatments in the gnotobiotic growth bags described above (*N* = 20 per treatment for B73 and Mo17; *N* = 15 per treatment for B73xMo17). Prior to planting, the kernels were weighed and distributed evenly to ensure that no systematic differences in seed size among the treatments. Bags were arranged into randomized, replicated blocks in a growth chamber in the Duke University Phytotron (12-hr days, 27^℃^C/23^℃^C, ambient humidity) and uprooted after one month of growth for measurement of shoot fresh weight and root fresh weight. Fisher’s Exact Test was used to compare germination proportions between genotypes within each treatment. We applied two-way ANOVA to linear mixed-effects models of biomass with Genotype, Treatment, and their interaction as fixed predictor variables, and Block as a random-intercept term. *F*-tests with Type III sums of squares were used for significance testing of fixed effects, and pairwise contrasts were performed using Tukey’s *post-hoc* procedure. Likelihood ratio tests were used for significance testing of random effects.

### Experiment 3

(September-November 2019). To determine whether MDH could be observed under field conditions, we conducted an on-farm soil sterilization experiment at the Central Crops Research Station. Total bed width was 152 cm furrow to furrow, and beds were 20 cm high with a 76 cm width at the top. Five treatments were established in a complete block design: steam-only (1 hr, 5 bar); allyl isothiocyanate (AITC; 280 L/ha); AITC (280 L/ha) followed by steam (1 hr, 5 bar); non-treated control; and chloropicrin (320 L/ha Pic-Clor 60). AITC and chloropicrin were applied September 11th, 2019 through shank application in raised beds. After fumigation, raised beds were covered with black Totally Impermeable Film (TIF) plastic. Steam was applied Sept. 27th using a SIOUX SF-25 Natural Gas Steam Generator (SIOUX Inc., Beresford, SD). The steam generator has a net heat input of 1.01e6 BTU/hr and an average steam output of 383 kg/hr. The steam generator was mounted on a flatbed trailer and connected to natural gas tanks, a 1,300 L water tank and a natural gas electrical generator (Supplementary Fig. 4f). Steam was applied consistently for 1 hr at 5 bar, injecting steam in 12 cm depth under TIF plastic using custom-made steam-graded spike hoses (Supplementary Fig. 4e). Temperature was monitored in different depths using HOBO U12 Outdoor/industrial data logger (Onset Computer Corporation, Bourne, MA). The maximum temperatures reached in the steam-only treatment were 100 °C in 12 cm depth; in the AITC + steam treatment, maximum temperatures of 66 °C were measured (Extended Data Fig. 4d). Kernels were hand-planted 4 cm deep into slits in the plastic (6” spacing between slits, with two seeds 3” apart on opposite ends of each slit), randomized within 4 blocks per treatment and 7 sub-blocks per block. To reduce seed-borne microbial load while in the field, we soaked kernels in 3% hydrogen peroxide for 2 minutes and rinsed in sterile diH2O immediately prior to planting. Plants were monitored for emergence three times (5, 8, and 12 days after planting) and height was measured twice (15 and 27 days after planting). After 27 days of growth, plants from three of the treatments (chloropicrin, AITC+steam, and control) were uprooted and oven-dried for measurement of root and shoot biomass. For the biomass data, we applied two-way ANOVA to linear mixed-effects models with Genotype, Treatment, and their interaction as fixed predictor variables, and Block and Sub-block as random-intercept terms. For the height and leaf number data, we applied three-way repeated-measures ANOVA with Genotype, Treatment, Date, and all interactions as fixed predictors, and Plant, Block, and Sub-block as random-intercept terms. *F*-tests with Type III sums of squares were used for significance testing of fixed effects, and pairwise contrasts were performed using Tukey’s *post-hoc* procedure. Likelihood ratio tests were used for significance testing of random effects.

### Fumigation effects on *Pythium* survival (Experiment 3)

To evaluate the efficacy of the steam, steam + AITC, and chloropicrin treatments on *Pythium ultimum* and fusarium wilt, mixed soils samples (6 inches deep) were taken post treatment. Five samples were combined to one mixed sample, taken from a 5 ft x 5 ft sample area. Four mixed soil samples were taken per treatment. The survival of *P. ultimum* was assessed using the wet plating method on semi-selective medium (34). Corn meal agar (Sigma-Aldrich, St. Louis, MO) was prepared (17 g⋅L^−1^) and sterilized. After sterilization, a nonionic detergent (Tween 20, Thermo Fisher Scientific, Waltham, MA) (1 mL⋅L^−1^) was added. Rose bengal (Thermo Fisher Scientific) (25 µg⋅mL^−1^), rifampicin (Thermo Fisher Scientific) (10 µg⋅mL^−1^), ampicillin (Sigma-Aldrich) (25 µg ·mL^−1^), pimaricin (Sigma-Aldrich) (5 µg⋅mL^−1^) and benomyl (Sigma-Aldrich) (40 µg⋅mL^−1^) were added to the agar at 50 °C. After cooling, 0.5 – 1 g of wet inoculum was spread (same dilution) on a total of five plates in three replicates. Plates were incubated at room temperature in the dark. *P. ultimum* colonies were counted during the first 2 d after plating. The mean propagules per gram soil were calculated for all three replicates (34).

### Fumigation effects on soil microbial community viability (Experiment 3)

To assess the effects of the fumigation treatments on the field soil, we collected soil samples weekly beginning immediately before planting and ending 4 weeks (56 time points) after planting, when plants were harvested. Four soil cores per week were collected from each treatment to a depth of 25 cm; each core was divided into subsamples taken from depths 3-5 cm and 17-20 cm and kept on ice for 4-5 h, then stored at 4°C overnight and used for bacterial counts the following day. Soil suspensions were prepared by mixing 1 g fresh soil in 9 ml of 0.95% NaCl. The suspension was then homogenized with a micro homogenizer (OMNI International, Inc., Kennesaw, GA, USA) at 12000 rpm for one 60 s cycle. After homogenization, serial dilutions were prepared up to 10^-5^. Samples were plated on R2A 1/10 and VxylG media (35) using the 6×6 drop plate method (36) for dilutions 10^-2^ to 10^-5^. Plates were incubated in the dark at 25°C for 3 weeks. Colony forming units (CFUs) were counted at 3, 7, 14, and 21 days; we determined that 14 days was the best time to count colonies and thus we used only that timepoint to calculate CFU per g soil.

### Fumigation effects on bacterial and fungal microbiomes (Experiment 3)

We used high-throughput amplicon sequencing to assess how the on-farm fumigation methods affected the bacterial and fungal communities in the soil at large. Bulk soil samples were collected from the treated blocks at three timepoints: one, four, and six weeks after the treatments were applied. After the final measurements of plant phenotype, the roots of a representative subsample of plants in the control, chloropicrin, and AITC+steam treatments were harvested for microbiome quantification. DNA was extracted from soil and root samples using the DNeasy PowerSoil kit (QIAGEN, Inc., Hilden, Germany) and used as a template for PCR amplification of the V4 region of the bacterial 16S rRNA gene and the fungal ITS1, following established protocols (11). The resulting 16S-v4 and ITS1 amplicons were then sequenced in parallel on the Illumina MiSeq platform (V2 chemistry, 250-bp PE reads) to census the bacterial and fungal components of the microbiome, respectively.

Established bioinformatic pipelines were used to quality-filter, denoise, and assign taxonomy to the raw sequence reads (11). Sequences that were derived from plants or that could not be identified at the kingdom level were discarded; samples with insufficient data (<500 bacterial reads or <500 fungal reads) were removed from the dataset. Finally, amplicon sequence variants (ASVs) that were not detected at least 10 times (for fungi) or 25 times (for bacteria) in at least 3 samples were removed from the dataset. The final bacterial dataset included 75 samples with a median of 30109 reads per sample, comprising 1307 ASVs. Sequencing depth was lower on average for fungi; as a result the fungal dataset included 57 samples with a median of 6466 reads per sample, comprising 122 ASVs. To reduce stochastic variation due to differences in sequencing depth, we applied the variance-stabilizing transformation (37) to the resulting ASV counts; additionally, we calculated the standardized, log-transformed sequencing depth for each sample to use as a nuisance variable.

To test whether fumigation treatments altered soil microbiome composition, we used permutational MANOVA to partition variance in the bacterial and fungal communities among several sources: sequencing depth, timepoint, treatment, and the interaction between timepoint and treatment. To test whether fumigation treatments were also detectable in plant-associated communities at the end of the experiment, we conducted another permutational MANOVA to partition root microbiome variation among sequencing depth, genotype, treatment, and the interaction between genotype and treatment.

### Statistical tests for changes in strength of heterosis

For all experiments, we performed permutation tests to assess whether the change in strength of heterosis between sterile and nonsterile treatments was statistically significant. First, we used the estimated marginal means from the linear models described above to calculate the midparent heterosis (MPH) for each trait in each treatment:

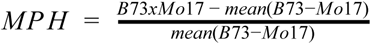

Second, we calculated “Δ_MPH_” as the difference in MPH between nonsterile and sterile treatments. Positive values of Δ_MPH_ indicate that heterosis was stronger in nonsterile conditions than in sterile conditions. Third, we re-calculated Δ_MPH_ for 999 datasets that had been permuted with respect to microbial Treatment, creating a distribution of Δ_MPH_ values that would be expected if Treatment had no effect on heterosis. Finally, we compared the observed Δ_MPH_ to this distribution using a one-tailed test of the null hypothesis that heterosis is not stronger in nonsterile conditions.

## Data and code availability

All raw data and original R code that support the findings of this study are freely available in a public repository (http://doi.org/10.5281/zenodo.4107065). Raw sequence reads are available in the NCBI SRA, BioProject #PRJNA669388.

## Author contributions

MRW, PBK, and MK conceptualized, designed, and supervised the experiments. CT performed soil viability assays and germination assays. FS, AB, MRW, and MK executed the synthetic community experiment. MRW executed the soil slurry experiment. MH, CT, FS, KC, SS, and PBK executed and/or collected data from the field experiment. MRW analyzed data and wrote the manuscript with input from all co-authors.

## Acknowledgements

We thank Dr. Saet-Byul Kim for suggesting the Whirl-Pak system; Dr. Ben Niu and Dr. Roberto Kolter for providing the SynCom strains; Hunter Smith for assistance with biomass measurements; Cathy Herring, Greg Marshall, Sophia Miller, Angie Mordant, Gitanjali Nanda Kafle, and the Central Crops Research Station staff for assistance with the field experiment; Dr. Carole Saravitz and the staff of the Phytotron at NCSU; Norman Hill, Greg Piotrowski, and the Duke University Phytotron staff. This material is based upon work supported by the National Science Foundation under Award No. OIA-1656006 and matching support from the State of Kansas through the Kansas Board of Regents. M.R.W. was also supported by a NSF National Plant Genome Initiative Postdoctoral Research Fellowship in Biology (IOS-1612951). Research in the laboratory of M.K. was supported by the NC State Chancellor’s Faculty Excellence Program Cluster on Microbiomes and Complex Microbial Communities, the USDA National Institute of Food and Agriculture, Hatch project 1014212, and grants from the Plant Soil Microbial Community Consortium and the Novo Nordisk Foundation (NNF19SA0059360 and NNF19SA0059362).

## Supplementary Information

**Supplementary Figure 1.**
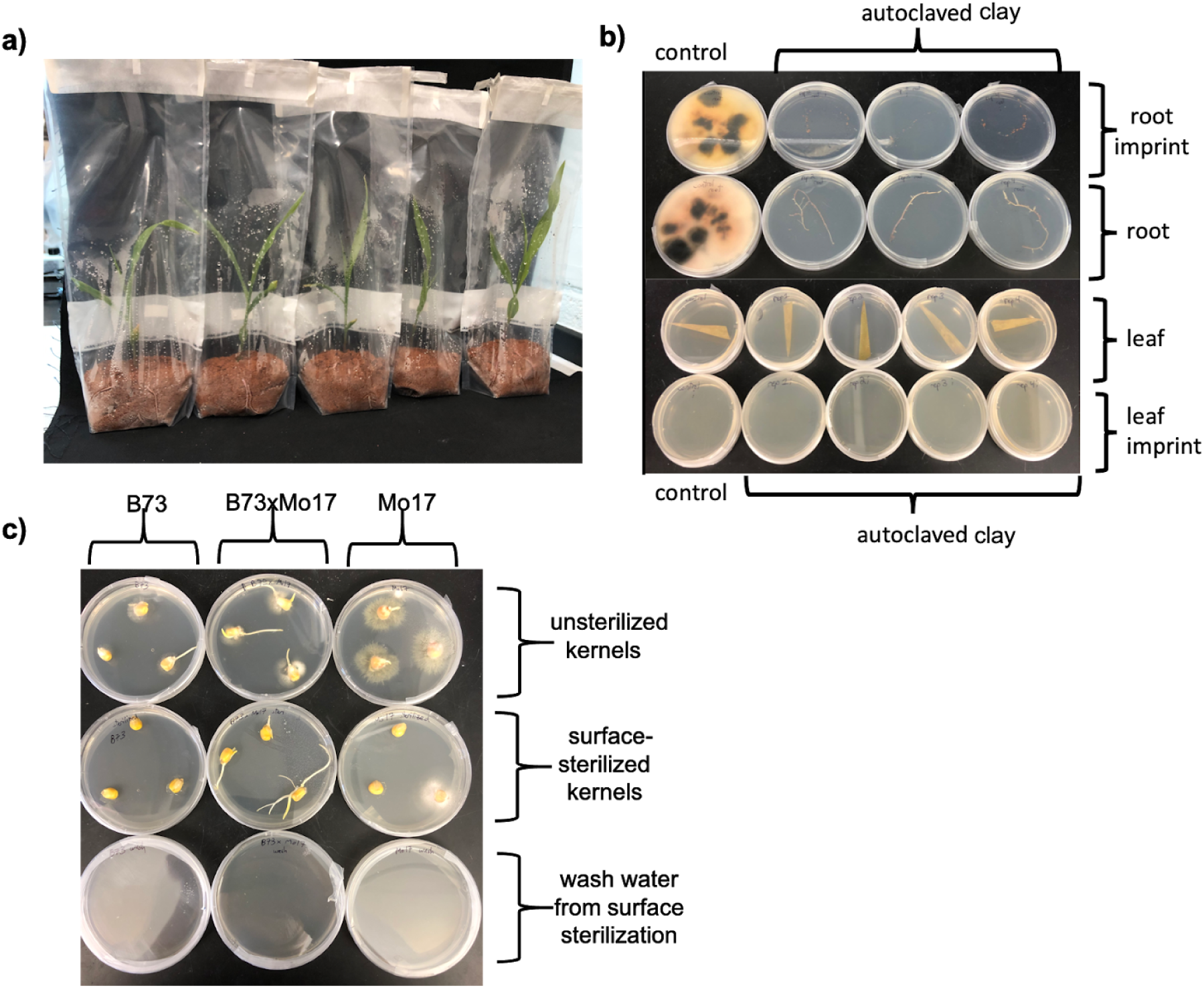
Gnotobiotic growth bags successfully created sterile conditions for maize plants up to 4 weeks old. **(a)** Kernels were planted into autoclaved calcined clay inside sterile Whirl-Pak bags and hydrated with sterile 0.5x MS. **(b)** Plating of roots and root imprints on potato dextrose agar (PDA) resulted in ample microbial growth for plants grown in non-autoclaved clay; in contrast, no growth was seen two weeks after plating for plants grown in gnotobiotic conditions. **(c)** Surface sterilization of kernels substantially reduced, but did not fully eliminate, seed-borne microbial load. Plating on malt extract agar (MEA) led to visible microbial growth from ~1 out of 9 surface-sterilized seeds (middle row). No growth was observed after plating the water from the final wash of seed surfaces (bottom row), indicating that this growth was most likely a seed endophyte that survived the surface sterilization.

**Supplementary Figure 2.**
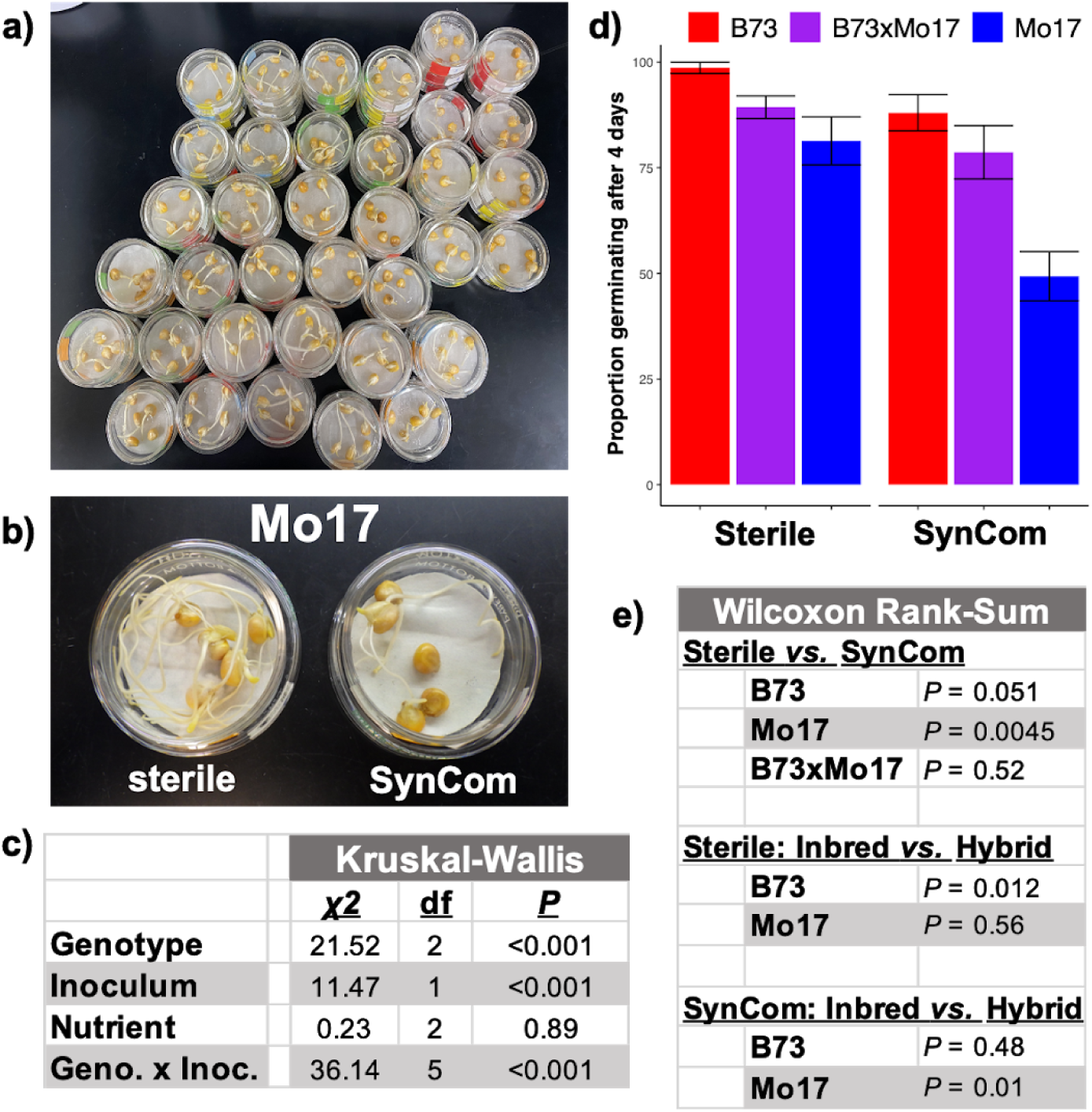
**(a)** Surface-sterilized kernels were plated on filter paper and hydrated with water, MS, or Hoagland’s solution either with or without the synthetic community (SynCom) of seven bacterial strains (*N =* 90 dishes; 5 kernels per dish). **(b)** Mo17 kernels in sterile water (left) or inoculated with the SynCom (right). **(c)** Kruskal-Wallis test demonstrated genotype-specific effects of the SynCom on germination. **(d)** Mean germination proportions (+/− 1 s.e.m.) are shown for each genotype after inoculation with the SynCom (right) or a sterile control (left). Results are averaged over all three nutrient treatments. **(e)** Wilcoxon rank-sum tests were used for pairwise comparisons of treatment-genotype combinations. *P-*values were adjusted to correct for multiple comparisons using the Benjamini-Hochberg method.

**Supplementary Figure 3.**
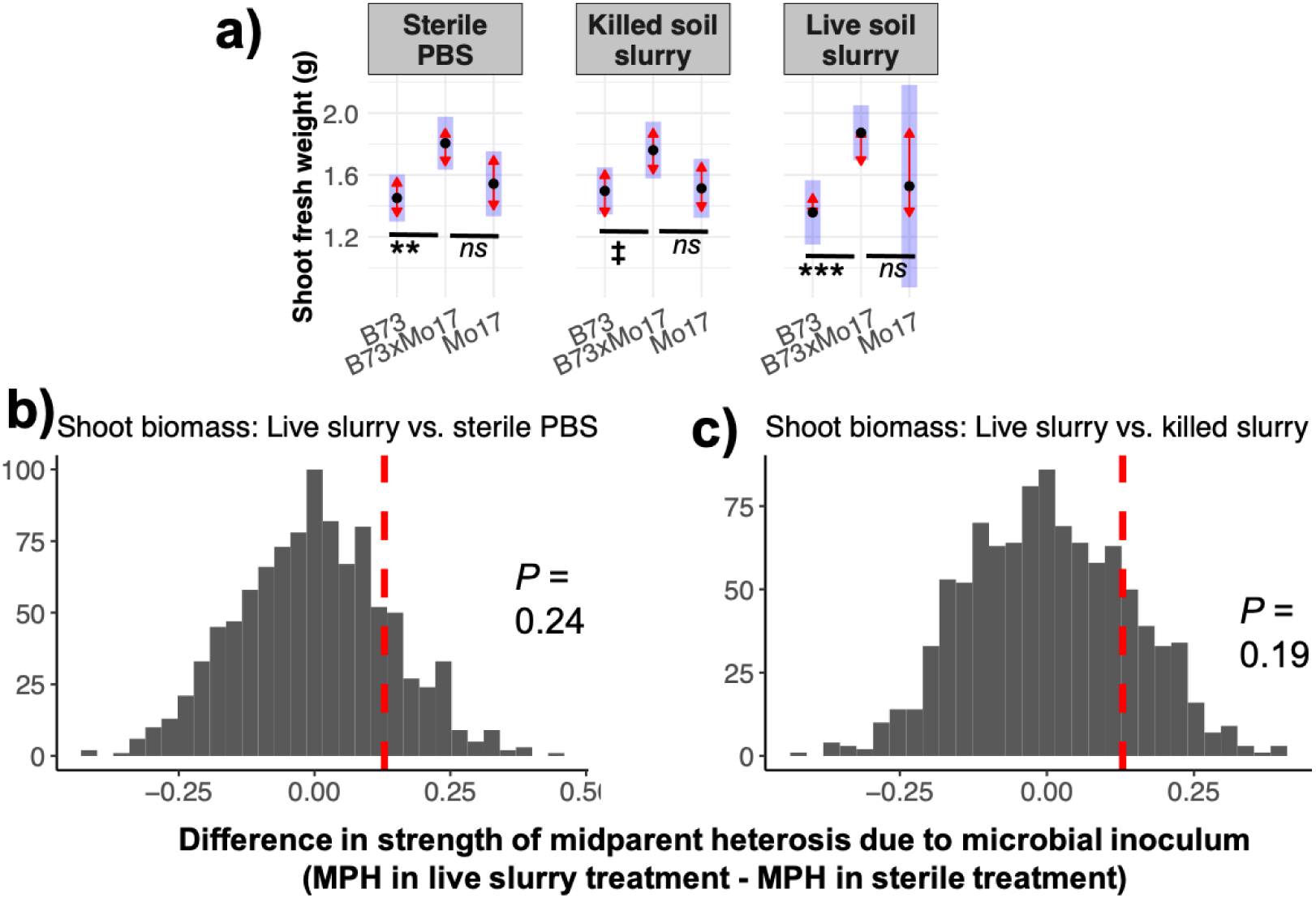
In experiment 2, maize kernels were grown in calcined clay inoculated with sterile PBS, a live soil slurry in PBS, or an autoclaved (killed) aliquot of the same soil slurry. Sterile conditions did not affect heterosis of shoot biomass. **(a)** A linear mixed-effects model was used for statistical inference of shoot biomass. Black points show the estimated marginal mean (EMM) trait values for each genotype in each treatment (values averaged over two timepoints); blue rectangles show the 95% CIs for these EMMs. The red arrows show the 95% CIs for pairwise tests between genotypes in each treatment after correction for the family-wise error rate using Tukey’s procedure; non-overlapping arrows indicate statistically significant differences (alpha=0.05). Detailed statistical results are provided in Supplementary Table 2. *N=*20 per inbred genotype per treatment, *N=*15 per hybrid per treatment. **^***^ *P*<0.001^;**^ *P*<0.01**; **^*^*P*<0.05**; **□ *P*<0.1**; ***ns P***>0.1 (Dunnett’s test of contrasts between each inbred line and the hybrid). **(c)** The strength of midparent heterosis (MPH) was calculated in each treatment using the EMM trait values. The observed difference in MPH between untreated control and fumigation treatments (∆MPH) was compared to the distributions of ∆MPH for 999 permutations of the data with respect to treatment, i.e., the distribution of ∆MPH if there were no effect of treatment. ***P***_∆MPH_ < 0.05 supports the alternate hypothesis that heterosis is stronger in untreated control soil than in fumigated soil.

**Supplementary Figure 4.**
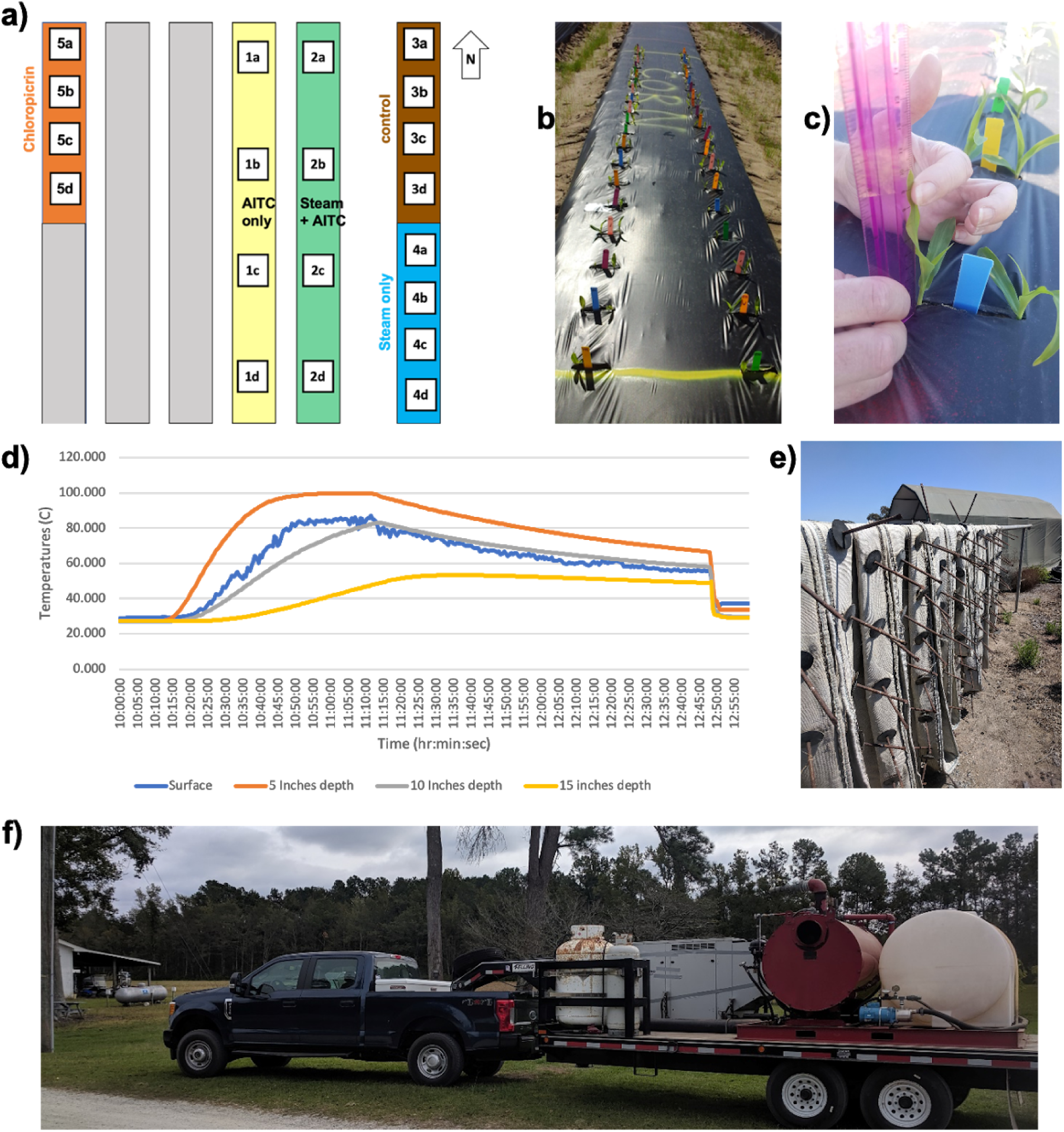
**(a)** Layout of field experiment (“Experiment 3”). Genotypes were randomized within 4 replicated blocks per soil treatment. **(b)** Maize seedlings emerging from a plastic-covered soil block. **(c)** Demonstration of height measurement for a seedling during the field experiment. **(d)** Temperature curve for the steam-only treatment. **(e)** Custom-made spike hoses used to inject steam into the soil. **(f)** SIOUX SF-25 natural gas steam generator with electrical generator, gas tanks, and water reservoir.

**Supplementary Figure 5.**
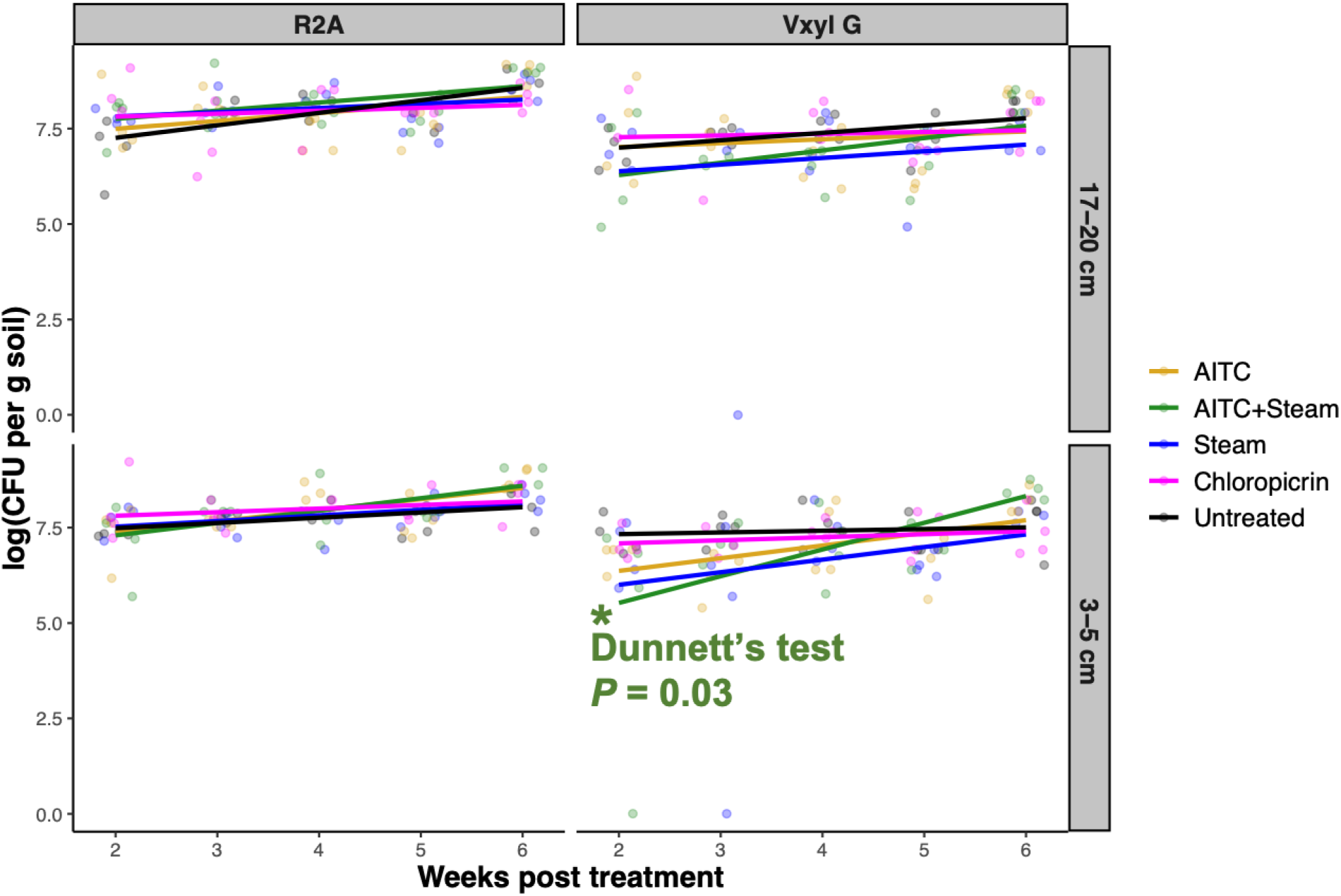
The density of viable bacteria (colony forming units g^−1^ soil) was measured at two depths in all treated and untreated field soils. Beginning two weeks after treatments were applied, four samples per treatment were taken weekly for the duration of the experiment. log(CFU g^−1^) was measured using two different microbial growth media (R2A and Vxyl G) and was modeled as a function of Treatment, Depth, Week (continuous variable), and all interactions. Slopes of log(CFU g^−1^) are shown for each treatment. Dunnett’s procedure was used to contrast the slope for each treatment to that of the untreated control; only the AITC + Steam treatment differed from the control at *P =* 0.05, and only in shallow soil when using Vxyl G media. *N* = 4 per treatment per depth per media per week.

**Supplementary Figure 6.**
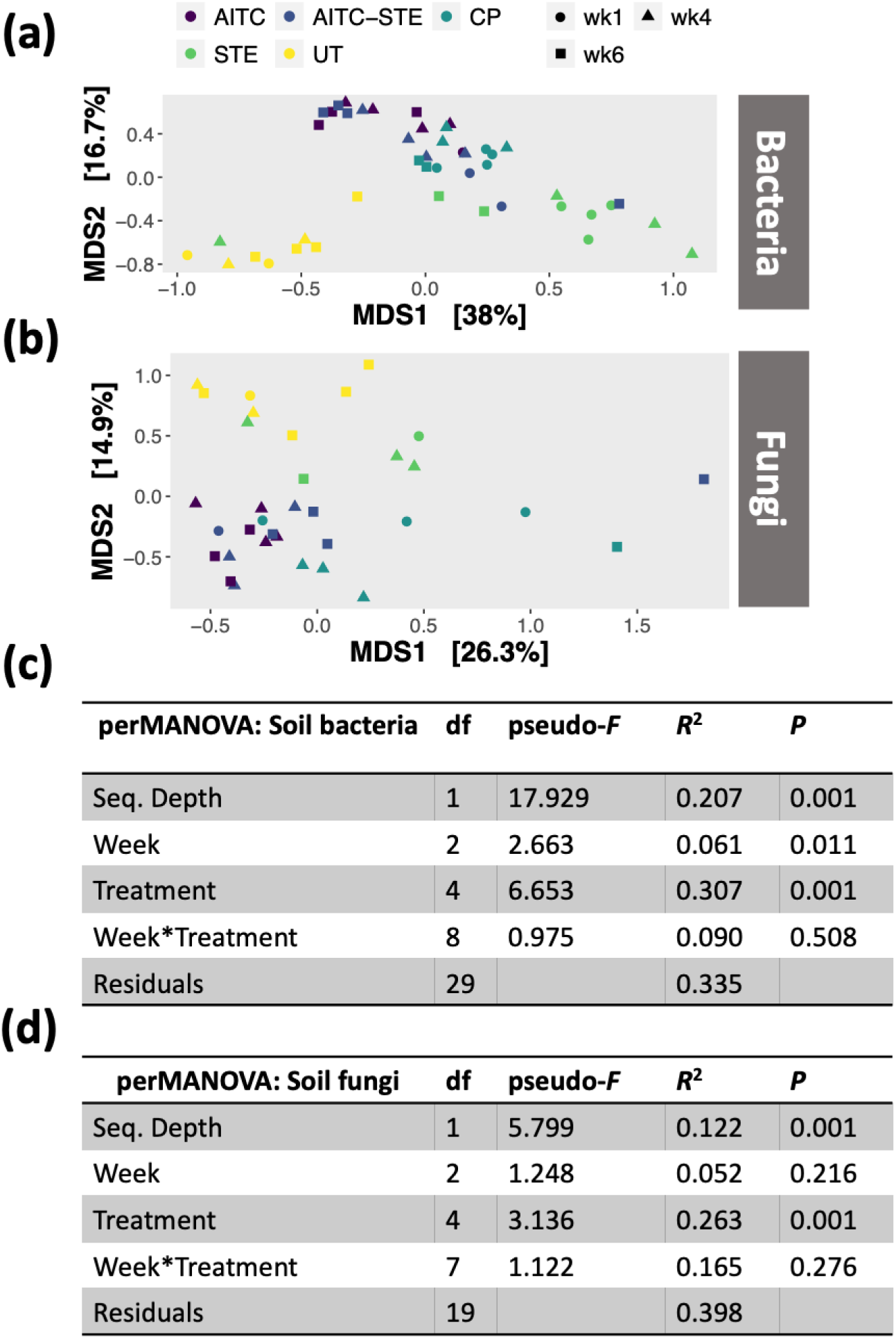
Soil fumigation treatments induced shifts in the composition of bacterial (a,c) and fungal (b,d) soil microbiomes that lasted at least 6 weeks. Plots (a-b) show unconstrained ordinations of Bray-Curtis distances between variance-stabilized ASV tables; the percentages on the axis labels show the proportion of community variation explained by each major axis. Effects of variation in sequencing depth were partialled out prior to ordination. Treatment abbreviations: AITC = allyl isothiocyanate; AITC-STE = allyl isothiocyanate + steam; CP = chloropicrin; STE = steam; UT = untreated control. Tables (c-d) show the results of permutational MANOVA to partition variance in bacterial and fungal communities among week, treatment, and the interaction between them after controlling for variation in sequencing depth.

**Supplementary Figure 7.**
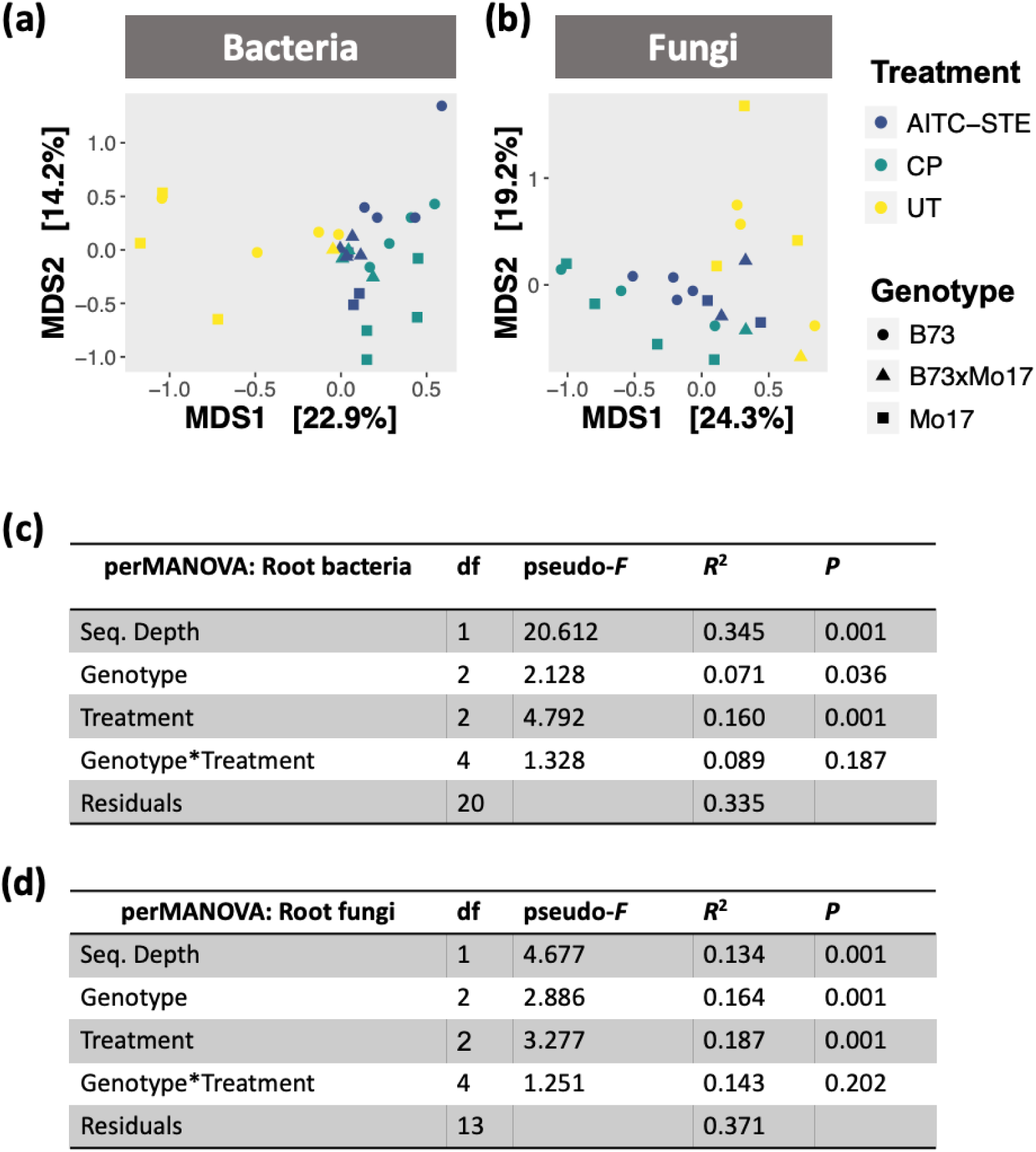
Bacterial (a,c) and fungal (b,d) microbiomes in maize roots harvested at the end of the field experiment were influenced by soil fumigation prior to planting. Plots (a,b) show unconstrained ordinations of Bray-Curtis distances between variance-stabilized ASV tables; the percentages on the axis labels show the proportion of community variation explained by each major axis. Effects of variation in sequencing depth were partialled out prior to ordination. Treatment abbreviations: AITC-STE = allyl isothiocyanate + steam; CP = chloropicrin; UT = untreated control. Tables (c,d) show the results of permutational MANOVA to partition variance in bacterial and fungal communities among host genotype, treatment, and the interaction between them after controlling for variation in sequencing depth.

**Supplementary Figure 8.**
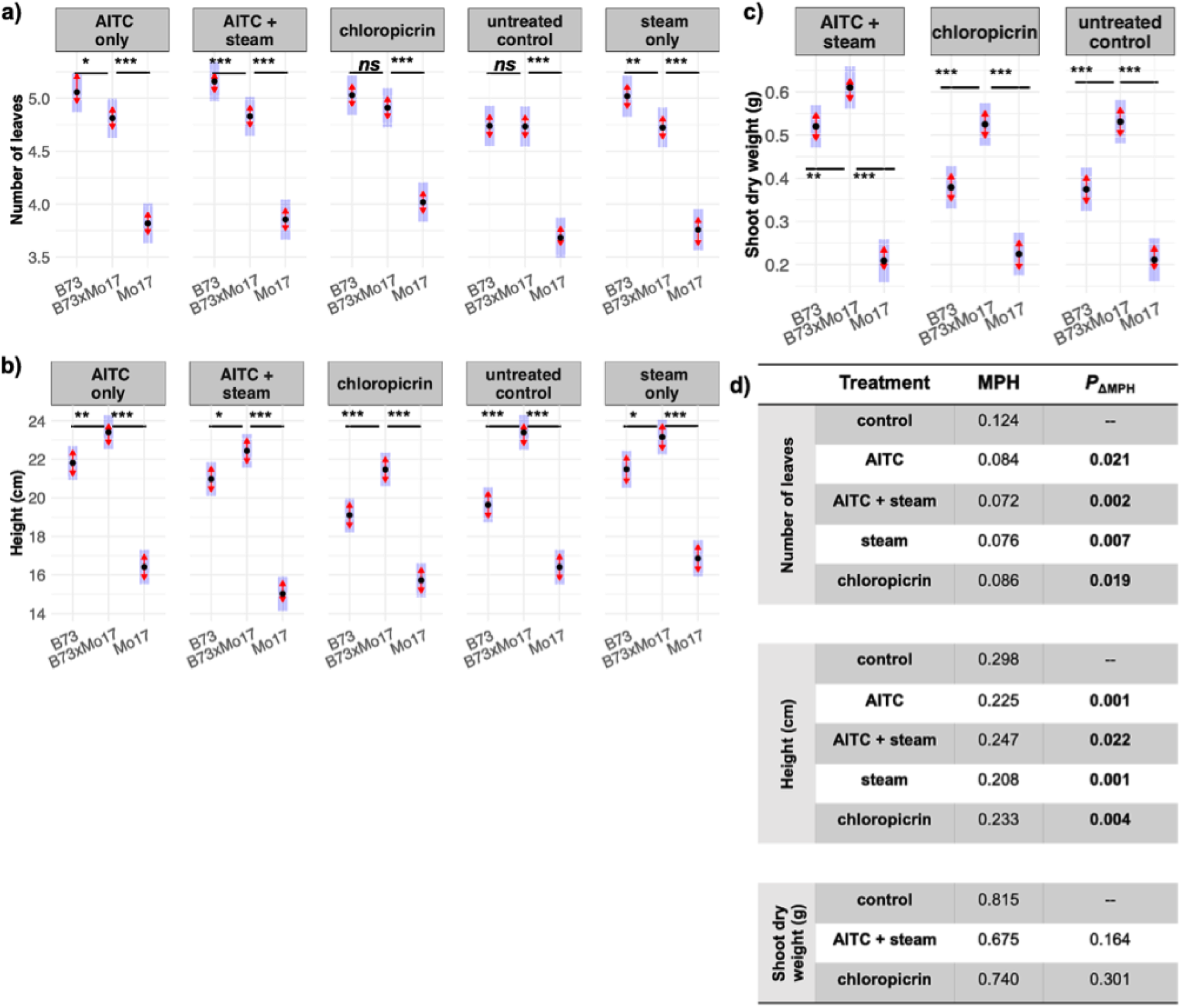
In Experiment 3, we grew maize in the field from seeds planted into untreated soil, soil fumigated with chloropicrin, or soil fumigated with AITC and/or steamed. **(a)** Leaf number and **(b)** height were measured for all treatments; shoot biomass **(c)** was measured for three treatments. Black points show the estimated marginal mean (EMM) trait values for each genotype in each treatment (values averaged over two timepoints); blue rectangles show the 95% CIs for these EMMs. The red arrows show the 95% CIs for pairwise tests between genotypes in each treatment after correction for the family-wise error rate using Tukey’s procedure; non-overlapping arrows indicate statistically significant differences (alpha=0.05). Detailed statistical results are provided in Supplementary Table 4. Effects on root and shoot biomass are presented in Figure 3. Effects on germination are presented in Supplementary Figure 9. **^***^ *P*<0.001**; **^**^ *P*<0.01**; **^*^*P*<0.05**; **□ *P*<0.1**; ***ns P***>0.1 (Dunnett’s test of contrasts between each inbred line and the hybrid) **(d)** The strength of midparent heterosis (MPH) was calculated for each trait in each treatment using the EMM trait values. The observed difference in MPH between untreated control and fumigation treatments (∆MPH) was compared to the distributions of ∆MPH for 999 permutations of the data with respect to treatment, i.e., the distribution of ∆MPH if there were no effect of treatment. ***P***_∆MPH_ < 0.05 supports the alternate hypothesis that heterosis is stronger in untreated control soil than in fumigated soil.

**Supplementary Figure 9.**
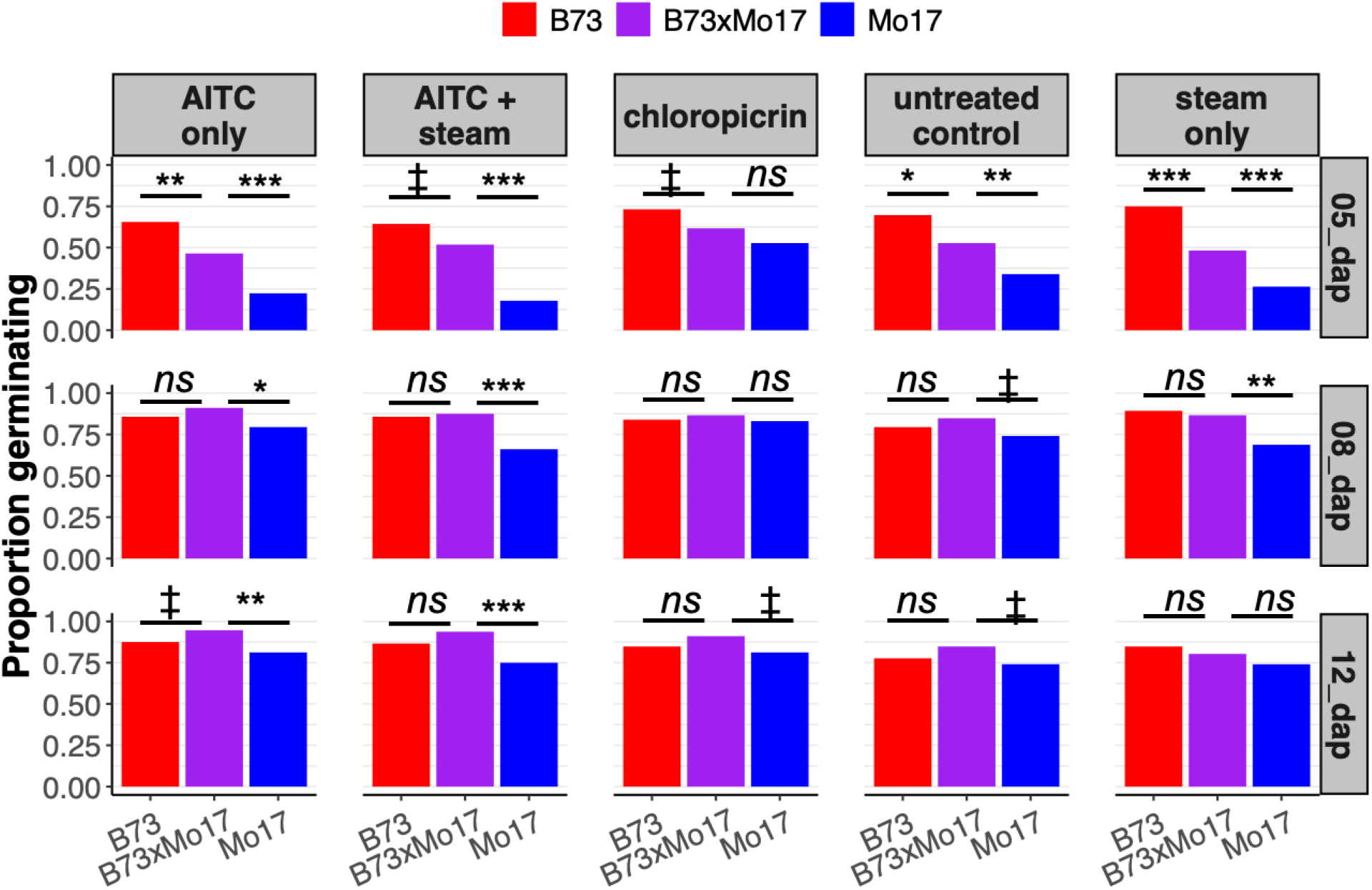
In Experiment 3, we grew maize in the field from seeds planted into untreated soil, soil fumigated with chloropicrin, or soil fumigated with AITC and/or steamed. Bar heights show the proportion of seeds that successfully germinated for each genotype in each treatment (columns) at each of three timepoints (rows; d.a.p. = days after planting). ***N =*** 112 per genotype per treatment. Statistical inference is from Fisher’s Exact Test. **^***^ *P*<0.001**; **^**^ *P*<0.01 ^;*^*P*<0.05**; **□ *P*<0.1**; ***ns P***>0.1

**Supplementary Table 1.** Analysis of variance (ANOVA) for linear models of fresh weight of **(a)** roots and **(b)** shoots in a gnotobiotic growth experiment, in which sterile maize seeds were inoculated with a synthetic community of seven bacterial strains or with a sterile buffer control (Fig. 1). *F*-tests with Type III sums of squares were used for significance testing.

|  | (a) | Root fresh weight | (b) | Shoot fresh weight |
| --- | --- | --- | --- | --- |
| **Genotype** | | $F_{2,33} = 1.67$<br>$P = 0.20$ | | $F_{2,33} = 0.25$<br>$P = 0.78$ |
| **Treatment** | | $F_{1,33} = 12.68$<br>$P = 0.0011$ | | $F_{1,33} = 2.67$<br>$P = 0.11$ |
| **Geno. x Treatment** | | $F_{2,33} = 1.34$<br>$P = 0.28$ | | $F_{2,33} = 0.79$<br>$P = 0.46$ |

**Supplementary Table 2.** Analysis of variance (ANOVA) for linear mixed models of fresh weights of maize seedling **(a)** roots and **(b)** shoots in in a gnotobiotic growth experiment, in which sterile maize seeds were inoculated with a live soil slurry, an autoclaved aliquot of the same slurry, or a sterile buffer control (Fig. 2). *F*-tests with Type III sums of squares were used for significance testing of fixed effects, and likelihood ratio tests were used for significance testing of the Block random effect. The Kenward-Roger method was used to estimate denominator degrees of freedom.

|  | (a) | Root fresh weight | (b) | Shoot fresh weight |
| --- | --- | --- | --- | --- |
| **Genotype** | | $F_{2,62} = 1.09$<br>$P = 0.34$ | | $F_{2,60} = 10.46$<br>$P = 0.00013$ |
| **Treatment** | | $F_{2,75} = 3.70$<br>$P = 0.029$ | | $F_{2,78} = 0.01$<br>$P = 0.98$ |
| **Geno. x Treatment** | | $F_{4,71} = 2.20$<br>$P = 0.078$ | | $F_{4,74} = 0.58$<br>$P = 0.68$ |
| **Block** | | $\chi^2_1 = 7.44$<br>$P = 0.0064$ | | $\chi^2_1 = 3.26$<br>$P = 0.071$ |

**Supplementary Table 3.** Analysis of variance (ANOVA) for *Pythium* sp. ppg (propagules per g soil) measured in treated and untreated field soils. ANOVA: P < 0.001; F = 17.01; df = 4. Fisher-LSD (p=0.05)

| | Mean ppg ± SEM | Fisher LSD ($P=0.05$) |
| --- | --- | --- |
| Untreated control | 3057.77 ± 512.1 | a |
| Steam | 1911.11 ± 644.9 | b |
| AITC | 180 ± 69.2 | c |
| AITC+Steam | 125 ± 57.2 | c |
| Chloropicrin | 44.4 ± 32.1 | c |

**Supplementary Table 4.**
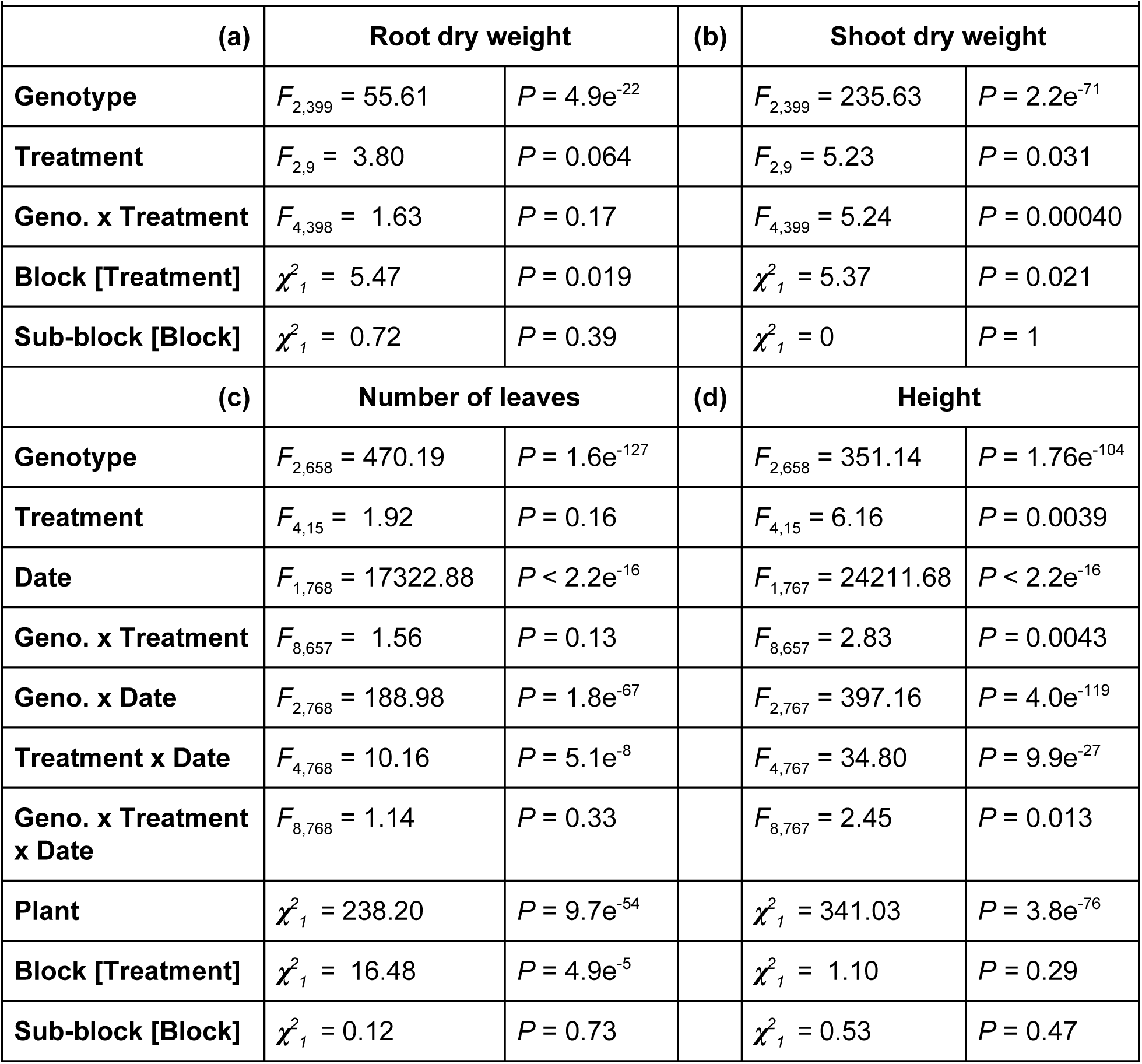
**(a-b)** Analysis of variance (ANOVA) for linear mixed models of dry weights of field-grown maize seedling roots and shoots measured at 27 days after planting (d.a.p.) **(c-d)** Repeated-measures ANOVA for linear mixed models of seedling height and leaf number measured at 15 d.a.p. and 27 d.a.p. *F*-tests with Type III sums of squares were used for significance testing of fixed effects, and likelihood ratio tests were used for significance testing of random effects. The Kenward-Roger method was used to estimate denominator degrees of freedom.

